# BioPipelines: Accessible Computational Protein and Ligand Design for Chemical Biologists

**DOI:** 10.64898/2026.03.11.711024

**Authors:** Gianluca Quargnali, Pablo Rivera-Fuentes

## Abstract

Deep learning methods for protein structure generation, sequence design, and structure and property prediction have created unprecedented opportunities for protein engineering and drug discovery. However, using these tools often requires navigating incompatible software environments, diverse input/output formats, and high-performance computing infrastructure, any of which may hinder adoption by primarily experimental chemical biology laboratories. Here we present BioPipelines, an open-source Python framework that allows researchers to define multi-step computational design workflows in a few lines of code. Additionally, its robust yet modular architecture provides a straightforward way to expand the toolkit with different functionalities, particularly by leveraging coding agents, with little effort. The framework currently integrates over 30 tools encompassing structure generation, sequence design, structure prediction, compound screening, and analysis. The same workflow code can be prototyped interactively in a Jupyter notebook and then submitted for production-scale runs without modification. We demonstrate applications in inverse folding, gene synthesis, de novo protein design, compound library screening, iterative binding site optimization, and fusion-protein linker optimization. We hope this framework will empower researchers, allowing them to focus on the scientific question rather than computational logistics. BioPipelines is available under the MIT license at https://github.com/locbp-uzh/biopipelines.

## Introduction

Over the past decade, and especially with the advent of deep learning models, protein engineering has shifted from complex, expert-oriented software suites like Rosetta^[1]^ to a scattered collection of tools designed to work in a “black-box” fashion with simple inputs. AlphaFold2,^[2]^ which takes only a protein sequence as input and yields a structural prediction in output, exemplifies this trend. Subsequent structure prediction methods, including AlphaFold3,^[3]^ RoseTTAFold3,^[4]^ Chai-1,^[5]^ and Boltz2^[6]^ operate similarly and have extended prediction to biomolecular complexes encompassing proteins, nucleic acids, and small molecules, with the latter also providing binding affinity estimation. Diffusion-based methods, including RFdiffusion,^[7]^ Chroma,^[8]^ and FrameDiff^[9]^ have made backbone generation routine, producing scaffolds with conditioned structural features. In parallel, inverse folding models, notably ProteinMPNN,^[10]^ have enabled rapid sequence design for target structures, with extensions such as LigandMPNN^[11]^ accommodating small-molecule binding partners. Dozens of other tools now exist, including tools for docking and protein-ligand interaction analysis, among others.

These advances are especially relevant to chemical biology, where researchers routinely need to engineer proteins with altered binding specificity, design enzyme variants for new substrates, or screen compound libraries against protein targets. However, a framework is lacking to systematically connect these tools in a simple, standardized way. A typical computational campaign might involve generating protein scaffolds around a ligand binding site, designing sequences compatible with that scaffold, predicting the structures of the resulting designs, and ranking them by predicted binding affinity. In practice, however, chaining these tools together presents significant challenges for laboratories without dedicated computational support. Each tool requires its own software environment, uses different input and output file formats, and must be configured for the computing cluster. Running a multi-tool workflow manually involves writing and debugging shell scripts, tracking intermediate files across tools, and managing job dependencies on the cluster scheduler. These logistical hurdles are often the rate-limiting step in adopting computational design.

Some workflow frameworks have been developed to address this problem. ColabFold pioneered the democratization of protein folding tools,^[12]^ but it has remained largely focused on structure prediction models. Ovo^[13]^ offers a web interface built on the Nextflow^[14]^ workflow engine but requires a database server and containerization infrastructure. ProteinDJ^[15]^ achieves efficient multi-GPU parallelism but restricts users to nine predefined pipeline configurations without support for custom workflows or iterative optimization. ProtFlow^[16]^ provides Python wrappers around design tools with cluster job management but the absence of typed data for common entities in chemical biology impairs its modularity. Moreover, it requires writing verbose configuration code and maintaining a running Python process throughout execution.

Here we present BioPipelines, a framework designed to make computational protein and ligand design accessible to research groups with minimal computational expertise. In BioPipelines, workflows are expressed in a compact, experiment-like syntax in which tools interact through standardized biomolecular data streams. Pipelines can be inspected and tested interactively, for example in Jupyter, and be submitted to a SLURM^[17]^ computing cluster without modification. Further, the framework can easily be extended to incorporate diverse tools, particularly by leveraging artificial intelligence (AI) coding agents. We demonstrate these capabilities through a series of application examples spanning common tasks in chemical biology.

## Results and Discussion

### Software design

BioPipelines was designed with three objectives in mind: 1) abstraction: pipelines read like a description of an experiment, and run with minimal user-side tuning of the underlying computational infrastructure; 2) modularity: arbitrary tools can be implemented following simple, biomolecular-informed rules, so that they can serve as both input and output to pre-existing tools according to a standardized interface; and 3) testability: pipelines can be tested in interactive environments like Jupyter notebooks, and intermediate results can be easily inspected. We briefly discuss the main implementation here before focusing on some usage case examples, and redirect the interested reader to the supporting information, where a technical description of the architecture is provided.

Abstraction is achieved by separating configuration and execution into two conceptually distinct and subsequent phases. The configuration phase is fully orchestrated in Python, the most common high-level programming language in biochemical computational tools. A workflow is written as a compact script defining which tools to run, with what parameters, and how their outputs connect. At this stage, the framework predicts the file system structure and outputs that will result after execution and generates self-contained bash scripts handling tools execution and interfacing. Combined with a straightforward and standardized syntax rooted in Python’s context manager, it allows users to write pipelines in seemingly natural language, as demonstrated in the coming examples. Only in the second phase, after the Python script has terminated, the ensemble of bash scripts is executed. Therefore, no long-running orchestrator is required during cluster execution. Generated scripts can be inspected, modified, or resubmitted independently, serving as both execution artifacts and documentation of what was run.

The modularity of BioPipelines stems from a standardized representation of the entities on which tools operate and which tools produce. We identified a set of three basic types, which are: 1) structures (2D/3D data: .pdb, .cif, .sdf, etc), 2) sequences (1D data: proteins, DNA, RNA), and 3) compounds (SMILES, CCD, etc). These constitute the primary item streams flowing between different tools in a pipeline, together with tabular data associated with them. Nearly all tools of interest for chemical biology applications operate on one or more of these and, provided some tool-specific parameters, produce another set. For example, ProteinMPNN takes in structures and yields sequences, AlphaFold2 takes in sequences and yields structures, and Boltz2 takes in sequences and compounds and yields structures. Any such tool can be integrated into BioPipelines by writing an associated Tool class that predicts its output from its input. To complete the picture, BioPipelines allows arbitrary definitions of streams. For example, Boltz2 generates MSA files when they are not provided, and these can be recycled in subsequent runs. BioPipelines provides standardized functions to handle streams and operations on associated data tables like filtering and sorting.

Finally, testability is greatly enhanced by the fact that the same pipeline code can execute in on-the-fly mode when running in a Jupyter or Google Colab notebook. BioPipelines auto-detects the interactive environment and executes each tool immediately upon instantiation, streaming the console output to the notebook. Given that tool outputs are standardized, tools have a default implemented visual representation in notebooks: structures streams are displayed as interactive MOL files, plots are displayed in line, and so on. Tool outputs from an already run pipeline can also be loaded into Jupyter and displayed, as shown afterwards.

In the following sections, we showcase a few selected applications of BioPipelines to demonstrate its current capabilities, the simplicity and modularity of the code, and how these capabilities can be easily extended to include new tools.

### Application 1: Re-designing the sequence of a protein

The sequence of proteins largely dictates their physical properties, such as stability and solubility. Hence, it is of interest to design mutants that preserve the fold of the original protein but may lead to improved physical properties. BioPipelines can be used to generate alternative sequences of a given protein. In the example pipeline (**Figure 1**),^[18]^ we generate alternative sequences for ubiquitin. After defining the computational resources needed for the job, the starting structure of the protein is loaded from the Protein Data Bank (PDB: 4LCD).^[19]^ The sequence is redesigned with the ProteinMPNN model trained on soluble proteins, and the resulting sequences are folded with AlphaFold2 for inspection. Finally, these sequences are passed through a DNA encoder to produce codon-optimized DNA sequences ready for synthesis, cloning, and expression in *E. coli*. The DNAEncoder tool uses organism-specific codon usage tables from CoCoPUTs,^[20]^ applying, as a default option, a thresholded weighted sampling to avoid rare codons while reducing gene repeats.

**Figure 1.**
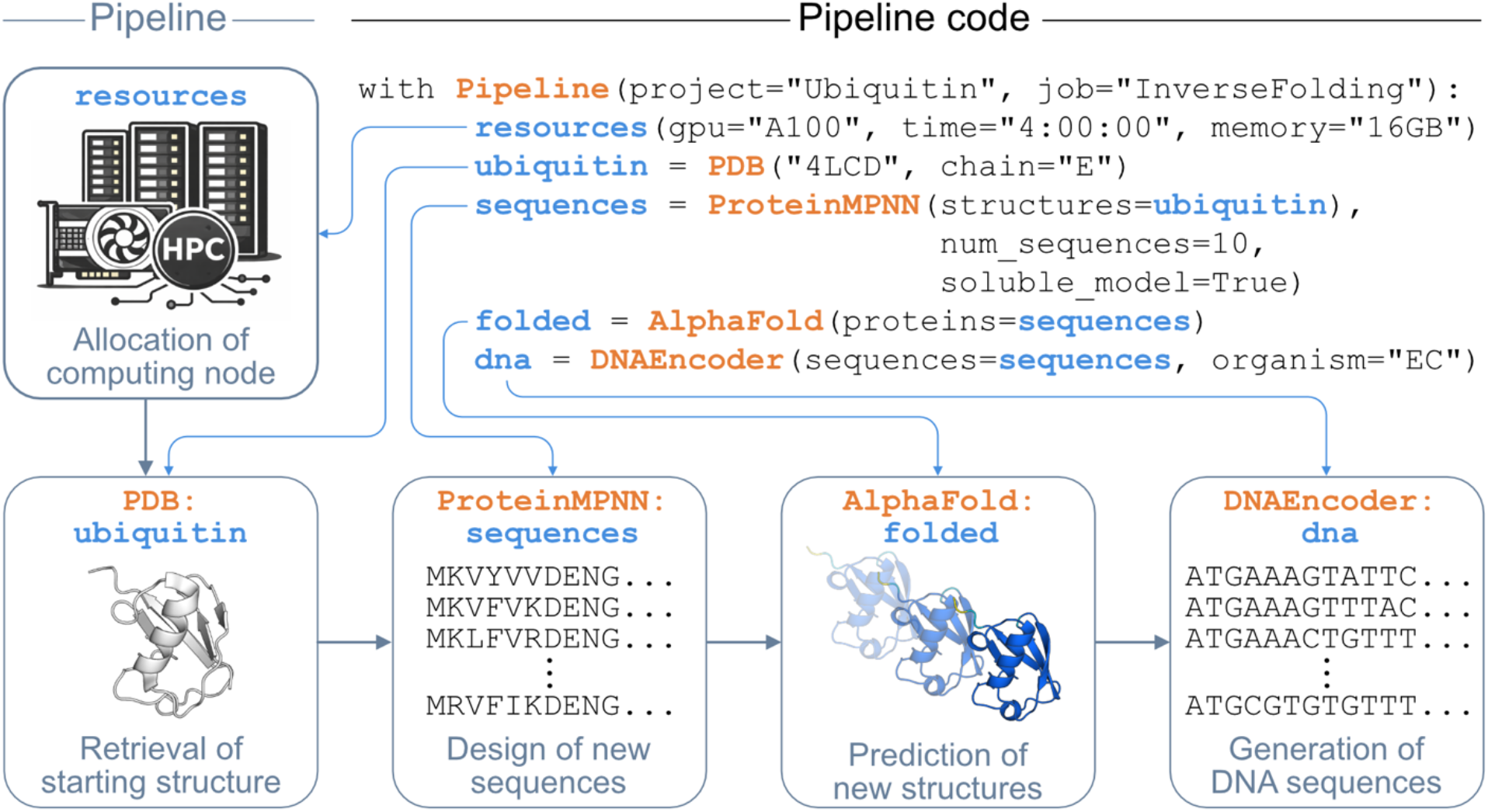
Pipeline and code for the design of alternative sequences of ubiquitin. BioPipelines’ Python classes are highlighted in orange and user-defined objects in blue.

Instead of relying on manual inspection of the structures, analysis and filtering within the pipeline are also supported. Commonly, one would discard folds that don’t resemble the original or that AlphaFold2 predicts with low confidence. This can easily be done with integrated tools in BioPipelines, by appending them to the workflow. For example, we can compute the RMSD of each new protein with respect to the parent protein and combine this information with a confidence metric from AlphaFold2, like the predicted local distance difference test (pLDDT), to obtain a refined set of sequences (**Figure S1**). Table operations, as well as ranking, filtering, and sorting based on tabular data, are handled by the BioPipelines tool Panda, which relies on the widespread Python package pandas.^[21]^

### Application 2: Redesigning a protein domain

Generative tools allow for designing entirely new proteins or parts of existing proteins, with countless potential applications in protein engineering, antibody design, and synthetic biology. BioPipelines can be used to seamlessly integrate de novo design of backbones, inverse folding, structure prediction, and analysis of results allowing the user to choose the specific model for each step of the design process. In the following example, the LID, a non-essential domain of the protein adenylate kinase (PDB: 4AKE), is redesigned. The pipeline is very similar to the previous example, starting with the allocation of computing resources and retrieval of the starting structure from the PDB (Figure 2). The only main difference is that RFdiffusion is called to generate ten new protein backbones, replacing the segment A118-160 (LID) with a new backbones of lengths between 50 and 70 amino acids. After this step, the workflow continues as for the ubiquitin case, with inverse folding of the new backbones by ProteinMPNN, which generates two sequences for each backbone generated by RFdiffusion, and structure validation by AlphaFold2.

**Figure 2.**
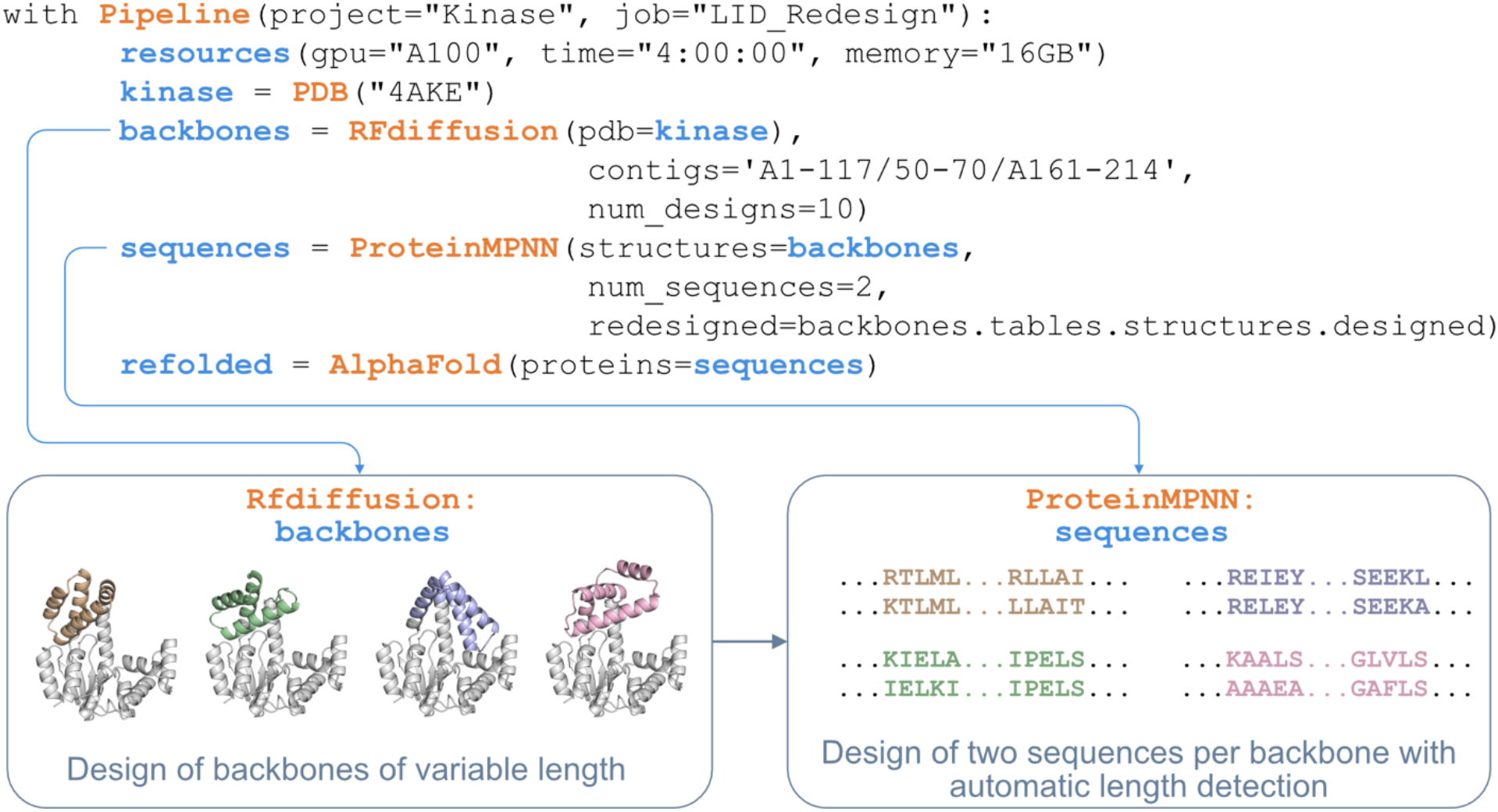
Biopipelines’ succinct implementation of the classic RFdiffusion-ProteinMPNN-AF2 pipeline. The resources allocation, starting structure retrieval (PDB class) and refolding (AlphaFold class) steps are equivalent to those depicted in Figure 1.

The `redesigned=backbones.tables.structures.designed` argument illustrates how information flows between tools. Each backbone has different designed positions, determined by contigs sampling of RFdiffusion at runtime, and this per-structure information is passed to ProteinMPNN without any manual file parsing, thus ensuring that ProteinMPNN designs sequences only for the newly generated RFdiffusion backbone, regardless of its length.

Pipelines involving RFdiffusion usually require hundreds, if not thousands, of designs to produce high-quality hits, depending on the objective. We can use the Panda tool again to not only filter the designs but also sort them based on any given metric produced and select the best ones. In **Figure S2** we show how to proceed from the pipeline to filter based on the conformational change of the non-designed portion of the protein, sort by model confidence, and generate a PyMOL session (.pse) file in which the top three structures are colored according to the AlphaFold coloring scheme for confidence in the designed regions, and white otherwise.

### Application 3: Screening a library of compounds against a target protein

Computational screening of compound libraries against protein targets is a common need in chemical biology, particularly for identifying lead compounds or optimizing binding interactions. BioPipelines provides tools for building combinatorial compound libraries and interface them with structure prediction tools. For example, Boltz2 can be used for cofolding of ligands with diverse biomolecules (proteins, DNA, RNA) and at the same time provides a state-of-the-art binding probability prediction (classification) and an associated—but arguably less reliable^[22]^—binding affinity (regression). In the following pipeline (**Figure 3**), a small library of tryptophan derivatives is screened against the homodimer of tryptophan repressor (TrpR) in the presence of its DNA operator (PDB: 1TRO).

**Figure 3.**
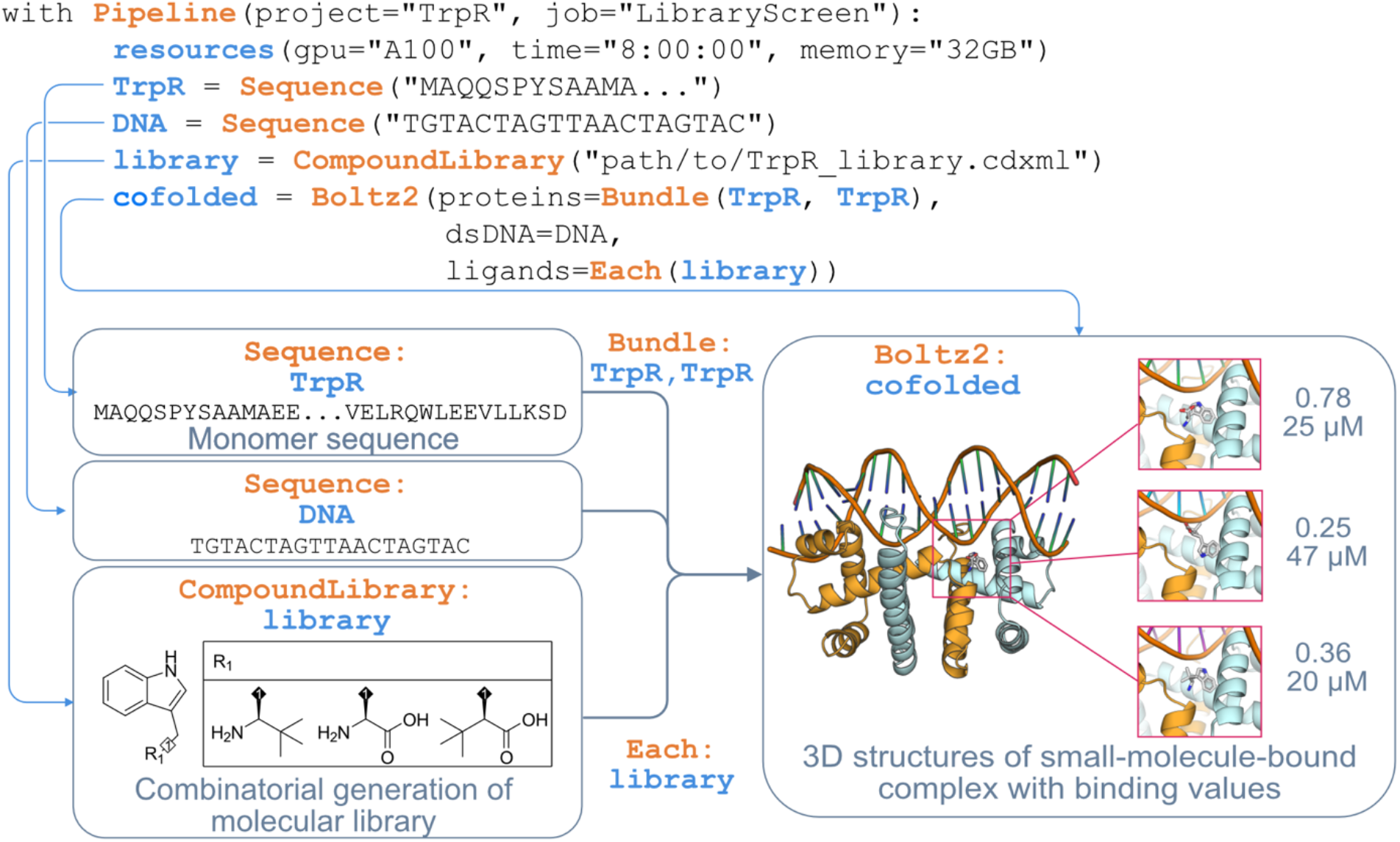
Definition of a branched compound library and interface with Boltz2. Bundle and Each control how to include entities into predictions: with Bundle on two copies of TrpR, we have both of them inside each prediction, resulting in a homodimer; with Each on the compound library, we have one prediction for each of the compounds. Affinities are provided as probability (unitless) and binding affinity (μM).

In BioPipelines, Boltz2 has been integrated into a simple syntax for declarative control over which entities and modifications to include while maintaining clarity through the combinatorial assistants Bundle and Each. In the above expression, we are instructing Boltz2 to cofold two copies of TrpR with each ligand present in the library, resulting in three separate predictions. This flexible syntax is maintained for more nested scenarios, for example, to calculate the affinity of each ligand in the presence of the homodimer and another molecule, or vice versa. The compound library can be generated combinatorially from either a ChemDraw file with appropriate R-group tables, or from a SMILES dictionary with declared branching points (**Figure S3**).

Finally, information flow allows us to easily gather useful metrics. In **Figure S4**, we use the Plot tool to plot the affinity predicted by Boltz2 against the R1 substituent from the library: the table “*compounds”* contains all the branching information for each compound. Importantly, the Plot tool generates a csv table containing the underlying data associated with each image file, so users can easily replot using their favorite software.

### Application 4: Modelling a Förster resonance energy transfer (FRET) calcium sensor

Genetically encoded FRET sensors typically sandwich a conformationally responsive sensing domain between donor and acceptor fluorescent proteins. A critical design question is which linkers keep both fluorescent domains properly folded while maximizing the change in FRET efficiency upon ligand binding. BioPipelines enables systematic screening of linker variants with structural validation under both apo and holo conditions. For example, the following pipeline (**Figure 4**) constructs a calmodulin-based calcium sensor by fusing EBFP (donor), calmodulin (sensing domain), and EYFP (acceptor) with variable-length flexible linkers. Both apo and Ca^2+^-bound forms are predicted with Boltz2, and the chromophore distance and inter-domain orientation are compared between states. A comprehensive screening, including both linker length and sequence, could easily be applied by using the Mutagenesis tool (**Figure S5**), and more structured linkers could be designed using RFdiffusion and ProteinMPNN. In the simple case shown here, the Fuse tool generates all linker-length combinations (4 x 4 = 16 constructs), concatenating the three domains with short flexible linkers. Both apo and holo forms are predicted for each construct, and multiple sequence alignments (MSAs) required for folding are recycled between the two, reducing impact on the server.

**Figure 4.**
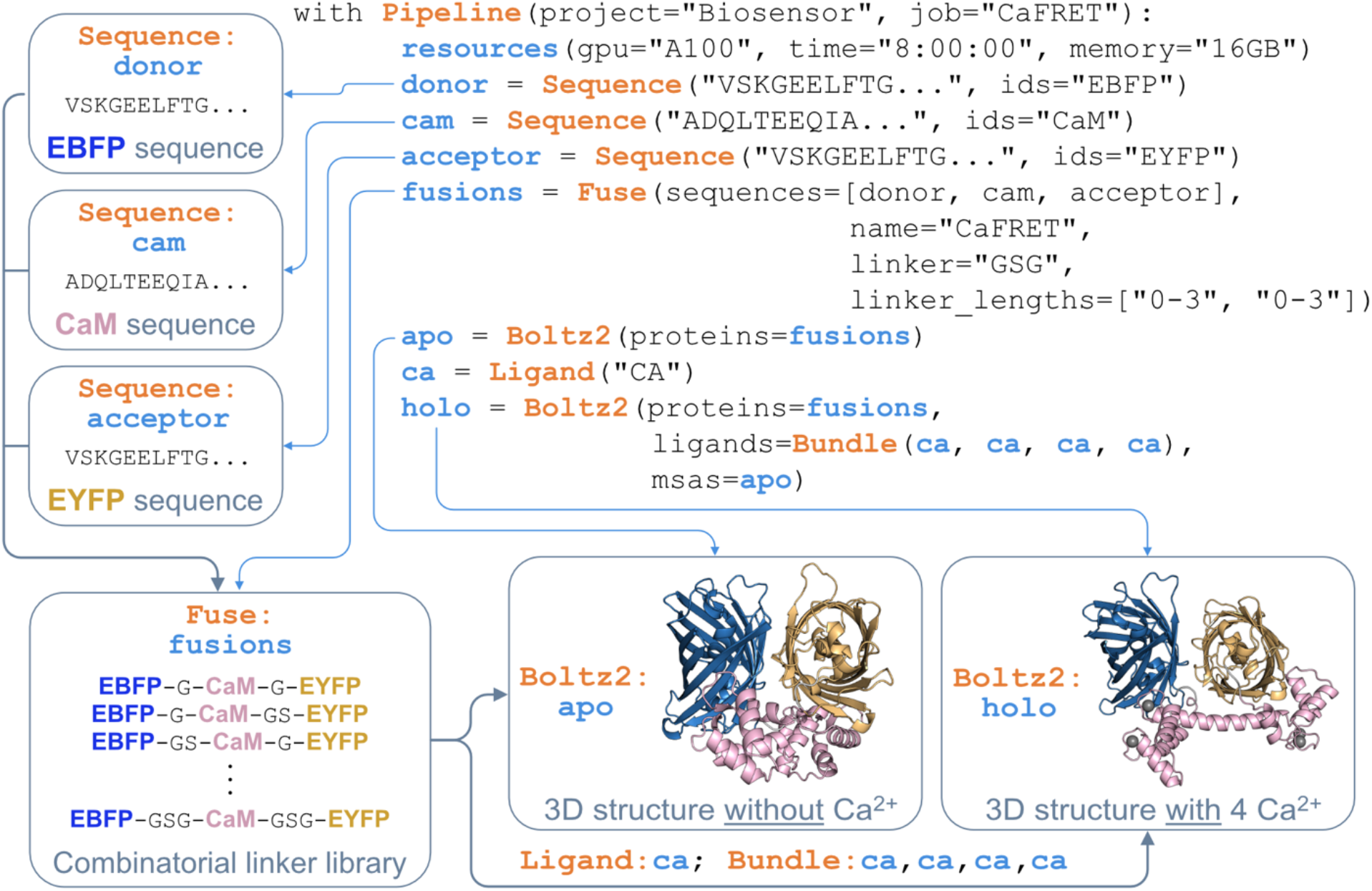
Modelling a FRET-based calcium sensor based on calmodulin and two fluorescent proteins. The server-intensive msas step is recycled for the holo structure determination from the apo calculations.

Further analysis of the designs could be carried out using the Distance tool (**Figure S6**) to measure the chromophore separation (residue 66 in the donor, residue -173 counting from the C-terminus in the acceptor), which is considered to be the most determinant geometric parameter for FRET efficiency when fluorophores are not rotationally restricted, an assumption we are making in this example. BioPipelines also provides the Angle tool for calculating angles and dihedrals, which could be used to extend the pipeline to measure inter-domain orientation. Finally, the Plot tool reveals which linker combinations produce the largest calcium-dependent change in FRET geometry, directly guiding sensor optimization.

### Beyond simple design pipelines: Iterative metric optimization

Many protein engineering campaigns can benefit from multiple rounds of design and evaluation, selecting a candidate for further engineering in the next iteration (e. g., directed evolution). With its modular nature, simple syntax, and tool-agnostic information flow, BioPipelines is a suitable platform to develop metric optimization methods. Our example pipeline demonstrates a simple case of cycles of optimization in a compact syntax, however, more sophisticated optimization schemes, such as machine-learning guided directed evolution,^[23,24]^ Bayesian optimization,^[25,26]^ active learning,^[27–29]^ or reinforcement learning,^[30]^ could also be implemented. In our simple pipeline (Figure 5), the objective is to find alternative sequences for the binding pocket of the periplasmic binding protein (PBP) NocT that bind non-covalently bound to histopine. The sequences are generated with LigandMPNN, profiled to obtain mutation frequencies, and the frequencies are used to generate 10 candidate mutants at each cycle based on weighted sampling. At each round, the structure and sequence giving the best predicted affinity are used as a template for the next iteration.

**Figure 5.**
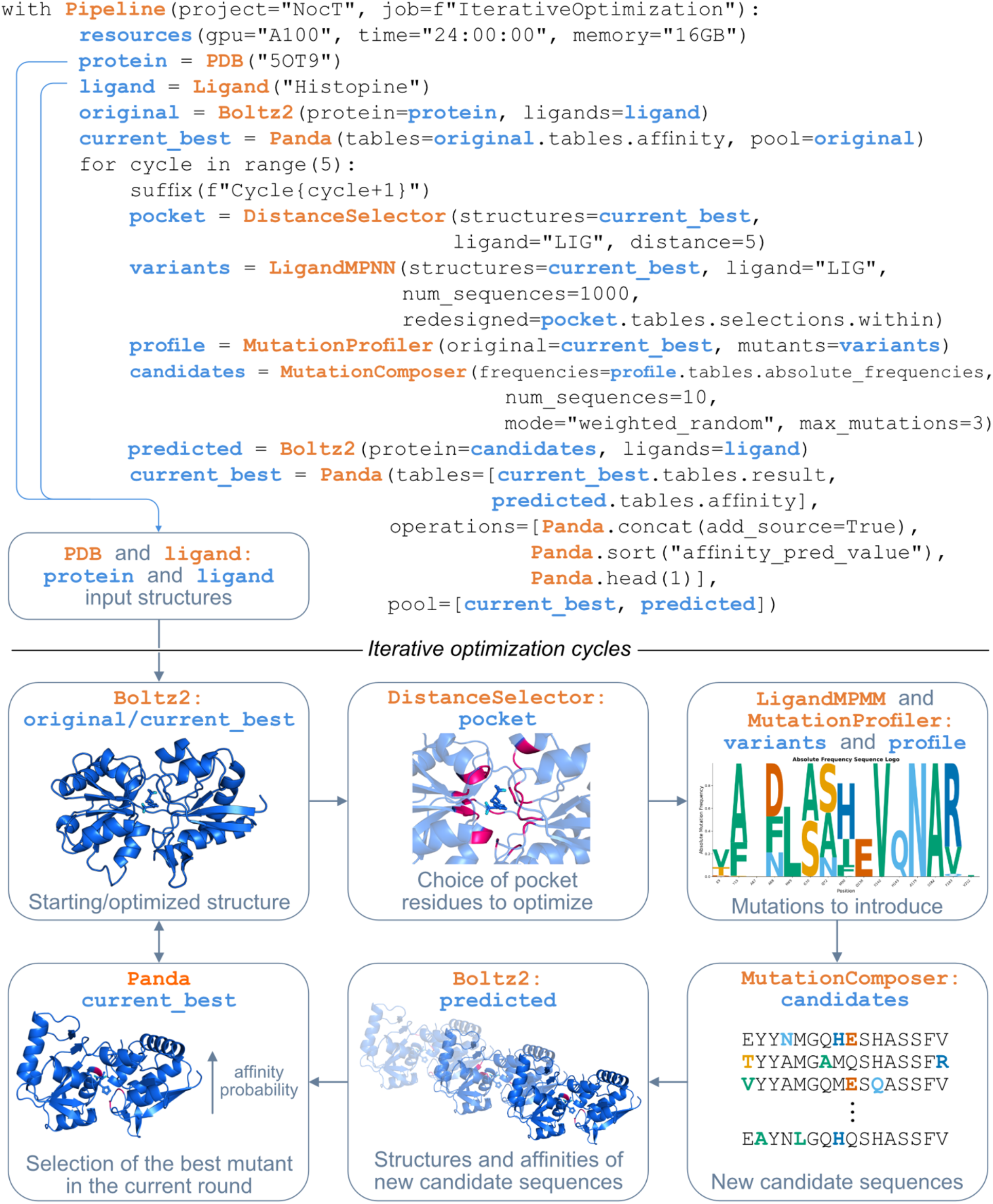
Computational evolution of a binding site based on computed affinity.

This example illustrates the potential of BioPipelines to develop new computational methods for metric optimization using any desired choice of protein generation, inverse folding, protein prediction, small-molecule generation, docking method, or structural analysis tool.

### Interactive prototyping

A key feature of BioPipelines is that the same workflow code can be tested prior to scaling up to inspect intermediate results, adjust parameters, and build the workflow step by step. Jupyter notebooks allow a rich visual representation of the output and are an ideal platform for this purpose. All tools that produce structures (e. g., AlphaFold, Boltz, RFdiffusion) automatically render an interactive py3Dmol 3D viewer. Images generated by the Plot tool are displayed inline together with the underlying data. These functionalities can be exploited in two ways, as illustrated in the Supporting Information, taking the second pipeline as an example (**Figure S7**): by running the pipeline directly in Jupyter, one can inspect intermediate results step by step as soon as execution is completed; and by using the Load tool, one can inspect any tool output from a previously run pipeline. Importantly, the loaded output has the same structure as the original one, thus behaves in the same way, allowing the user to employ it as input in downstream tools.

### Writing workflows and extending the framework

The motivation behind developing BioPipelines is that users with minimal coding experience can use and write custom workflows with ease. Because BioPipelines is a pure Python package with typed interfaces, users benefit from modern integrated development environment (IDE) features when writing workflows. Most modern IDEs (e. g., Visual Studio Code with Python and Pylance extensions) provide real-time autocompletion, parameter hints, and inline documentation for every tool and parameter. For example, typing “ `RFdiffusion` (“ immediately displays all available arguments with their types and descriptions, and passing an incorrect type triggers an underline warning before the script is ever run.

The standardized tool interface also makes it straightforward to extend BioPipelines with new tools using AI coding assistants. In practice, we have found that tools corresponding to published GitHub repositories can be implemented by providing Claude Code (Anthropic, Opus 4.6 model) with the repository URL and a single instruction: *“Implement a BioPipelines tool for the repository: <url> conforming to the existing tool standards*.*”* The assistant reads the repository’s installation instructions, input/output formats, and command-line interface, then generates a complete tool module, including the installation script, parameter validation, bash script generation, and output parsing that integrates directly into the framework. A representative example of this process is provided in the Supporting Information. This approach dramatically reduces the effort required to keep the framework current with the rapidly evolving landscape of computational biology tools and enables individual laboratories to add specialized or in-house tools without deep familiarity with the framework internals.

## Conclusions

BioPipelines makes computational protein and ligand design accessible to chemical biology laboratories by handling the computational logistics: environment management, file format conversion, cluster job scheduling, and data tracking that would otherwise require dedicated bioinformatics support. Researchers define workflows as concise Python scripts that read as descriptions of the scientific experiment, prototype interactively in Jupyter notebooks, and submit the same code for production runs. The framework’s support for compound library screening, iterative optimization, fusion protein engineering, and end-to-end gene synthesis preparation addresses common needs in chemical biology. BioPipelines currently integrates over 30 tools which we have organized by function in **Table S1**. New tools can be added by implementing four methods, outlined in the Supporting Information, and the framework provides base classes that handle environment activation, completion tracking, and standardized output formatting. The codebase is written to be easily understood by AI coding agents, enabling nonexperts to program new pipelines or add more specialized tools with minimal effort.

BioPipelines is freely available under the MIT license at https://github.com/locbp-uzh/biopipelines, with additional example pipelines. The full documentation can be found at https://biopipelines.readthedocs.io.

## Associated Content

### Supporting Information

Supplementary figures. Technical architecture details. Transcript of an AI-assisted tool implementation session, showing how a new tool was added to BioPipelines from a GitHub repository URL using Claude Code (Opus 4.6).

## Acknowledgments

We thank all members of the Laboratory of Chemical and Biological Probes at University of Zurich for valuable feedback. All computational work was carried out in the ScienceCluster (S3IT) of the University of Zurich. Substantial parts of the code were written or refactored by Claude Code (Opus 4.6).

## Funding

This work was supported by the Swiss National Science Foundation through project grant 200021-236456.

## Notes

https://github.com/locbp-uzh/biopipelines

https://biopipelines.readthedocs.io

### Competing Interest Statement

The authors have declared no competing interest.

## References

[1] K. T. Simons, R. Bonneau, I. Ruczinski, D. Baker, “Ab initio protein structure prediction of CASP III targets using ROSETTA” Proteins Struct. Funct. Genet. 1999, 37, 171–176.

[2] J. Jumper, R. Evans, A. Pritzel, T. Green, M. Figurnov, O. Ronneberger, K. Tunyasuvunakool, R. Bates, A. Žídek, A. Potapenko, A. Bridgland, C. Meyer, S. A. A. Kohl, A. J. Ballard, A. Cowie, B. Romera-Paredes, S. Nikolov, R. Jain, J. Adler, T. Back, S. Petersen, D. Reiman, E. Clancy, M. Zielinski, M. Steinegger, M. Pacholska, T. Berghammer, S. Bodenstein, D. Silver, O. Vinyals, A. W. Senior, K. Kavukcuoglu, P. Kohli, D. Hassabis, “Highly accurate protein structure prediction with AlphaFold” Nature 2021, 596, 583–589.

[3] J. Abramson, J. Adler, J. Dunger, R. Evans, T. Green, A. Pritzel, O. Ronneberger, L. Willmore, A. J. Ballard, J. Bambrick, S. W. Bodenstein, D. A. Evans, C.-C. Hung, M. O’Neill, D. Reiman, K. Tunyasuvunakool, Z. Wu, A. Žemgulytė, E. Arvaniti, C. Beattie, O. Bertolli, A. Bridgland, A. Cherepanov, M. Congreve, A. I. Cowen-Rivers, A. Cowie, M. Figurnov, F. B. Fuchs, H. Gladman, R. Jain, Y. A. Khan, C. M. R. Low, K. Perlin, A. Potapenko, P. Savy, S. Singh, A. Stecula, A. Thillaisundaram, C. Tong, S. Yakneen, E. D. Zhong, M. Zielinski, A. Žídek, V. Bapst, P. Kohli, M. Jaderberg, D. Hassabis, J. M. Jumper, “Accurate structure prediction of biomolecular interactions with AlphaFold 3” Nature 2024, 630, 493–500.

[4] N. Corley, S. Mathis, R. Krishna, M. S. Bauer, T. R. Thompson, W. Ahern, M. W. Kazman, R. I. Brent, K. Didi, A. Kubaney, L. McHugh, A. Nagle, A. Favor, M. Kshirsagar, P. Sturmfels, Y. Li, J. Butcher, B. Qiang, L. L. Schaaf, R. Mitra, K. Campbell, O. Zhang, R. Weissman, I. R. Humphreys, Q. Cong, J. Funk, S. Sonthalia, P. Liò, D. Baker, F. DiMaio, “Accelerating Biomolecular Modeling with AtomWorks and RF3”, bioRxiv 2025, 2025.08.14.670328

[5] Chai Discovery, J. Boitreaud, J. Dent, M. McPartlon, J. Meier, V. Reis, A. Rogozhnikov, K. Wu, “Chai-1: Decoding the molecular interactions of life”, bioRxiv 2024, 2024.10.10.615955

[6] S. Passaro, G. Corso, J. Wohlwend, M. Reveiz, S. Thaler, V. R. Somnath, N. Getz, T. Portnoi, J. Roy, H. Stark, D. Kwabi-Addo, D. Beaini, T. Jaakkola, R. Barzilay, “Boltz-2: Towards accurate and efficient binding affinity prediction” bioRxiv 2025, 2025.06.14.659707.

[7] J. L. Watson, D. Juergens, N. R. Bennett, B. L. Trippe, J. Yim, H. E. Eisenach, W. Ahern, A. J. Borst, R. J. Ragotte, L. F. Milles, B. I. M. Wicky, N. Hanikel, S. J. Pellock, A. Courbet, W. Sheffler, J. Wang, P. Venkatesh, I. Sappington, S. V. Torres, A. Lauko, V. De Bortoli, E. Mathieu, S. Ovchinnikov, R. Barzilay, T. S. Jaakkola, F. DiMaio, M. Baek, D. Baker, “De novo design of protein structure and function with RFdiffusion” Nature 2023, 620, 1089– 1100.

[8] J. B. Ingraham, M. Baranov, Z. Costello, K. W. Barber, W. Wang, A. Ismail, V. Frappier, D. M. Lord, C. Ng-Thow-Hing, E. R. Van Vlack, S. Tie, V. Xue, S. C. Cowles, A. Leung, J. V. Rodrigues, C. L. Morales-Perez, A. M. Ayoub, R. Green, K. Puentes, F. Oplinger, N. V. Panwar, F. Obermeyer, A. R. Root, A. L. Beam, F. J. Poelwijk, G. Grigoryan, “Illuminating protein space with a programmable generative model” Nature 2023, 623, 1070–1078.

[9] J. Yim, B. L. Trippe, V. D. Bortoli, E. Mathieu, A. Doucet, R. Barzilay, T. Jaakkola, “Fast protein backbone generation with SE(3) flow matching”, arXiv 2023, 2310.05297.

[10] J. Dauparas, I. Anishchenko, N. Bennett, H. Bai, R. J. Ragotte, L. F. Milles, B. I. M. Wicky, A. Courbet, R. J. de Haas, N. Bethel, P. J. Y. Leung, T. F. Huddy, S. Pellock, D. Tischer, F. Chan, B. Koepnick, H. Nguyen, A. Kang, B. Sankaran, A. K. Bera, N. P. King, D. Baker, “Robust deep learning–based protein sequence design using ProteinMPNN” Science 2022, 378, 49–56.

[11] J. Dauparas, G. R. Lee, R. Pecoraro, L. An, I. Anishchenko, C. Glasscock, D. Baker, “Atomic context-conditioned protein sequence design using LigandMPNN” Nat. Methods 2025, 22, 717–723.

[12] M. Mirdita, K. Schütze, Y. Moriwaki, L. Heo, S. Ovchinnikov, M. Steinegger, “ColabFold: making protein folding accessible to all” Nat. Methods 2022, 19, 679–682.

[13] B. Danny, “Ovo, an Open-Source Ecosystem for De Novo Protein Design”, bioRxiv 2025, 2025.11.27.691041

[14] P. Di Tommaso, M. Chatzou, E. W. Floden, P. P. Barja, E. Palumbo, C. Notredame, “Nextflow enables reproducible computational workflows” Nat. Biotechnol. 2017, 35, 316– 319.

[15] D. Silke, J. Iskander, J. Pan, A. P. Thompson, A. T. Papenfuss, I. S. Lucet, J. M. Hardy, “PROTEINDJ: A high-performance and modular protein design pipeline” Protein Sci. 2026, 35, e70464.

[16] Braun, M., & Tripp, A. (2024). “ProtFlow: A Python package to manage protein design workflows on computing clusters and local machines (v0.1.0)” [Software]. GitHub. https://github.com/mabr3112/ProtFlow

[17] A. B. Yoo, M. A. Jette, M. Grondona in Job Sched. Strateg. Parallel Process. (Eds.: D. Feitelson, L. Rudolph, U. Schwiegelshohn), Springer Berlin Heidelberg, Berlin, Heidelberg, 2003, pp. 44–60.

[18] L.-Y. Chen, W.-L. Lu, T. Pathania, I.-H. Chu, M.-R. Ho, W.-C. Chuang, Y.-C. Lou, T. I. Hung, Y. Miyanoiri, C. A. Chang, K.-P. Wu, “Mesostructured Water Enhances Stability of ProteinMPNN-Designed Ubiquitin-Fold Proteins” J. Am. Chem. Soc. 2026, jacs.5c19875.

[19] H. M. Berman, “The Protein Data Bank” Nucleic Acids Res. 2000, 28, 235–242.

[20] A. Alexaki, J. Kames, D. D. Holcomb, J. Athey, L. V. Santana-Quintero, P. V. N. Lam, N. Hamasaki-Katagiri, E. Osipova, V. Simonyan, H. Bar, A. A. Komar, C. Kimchi-Sarfaty, “Codon and Codon-Pair Usage Tables (CoCoPUTs): Facilitating Genetic Variation Analyses and Recombinant Gene Design” J. Mol. Biol. 2019, 431, 2434–2441.

[21] The pandas development team 2026, DOI 10.5281/ZENODO.3509134.

[22] G. Bret, F. Sindt, D. Rognan, “Assessing Boltz-2 Performance for the Binding Classification of Docking Hits” J. Chem. Inf. Model. 2026, 66, 1511–1521.

[23] Z. Wu, S. B. J. Kan, R. D. Lewis, B. J. Wittmann, F. H. Arnold, “Machine learning-assisted directed protein evolution with combinatorial libraries” Proc. Natl. Acad. Sci. 2019, 116, 8852–8858.

[24] S. Biswas, G. Khimulya, E. C. Alley, K. M. Esvelt, G. M. Church, “Low-N protein engineering with data-efficient deep learning” Nat. Methods 2021, 18, 389–396.

[25] A. Khan, A. I. Cowen-Rivers, A. Grosnit, D.-G.-X. Deik, P. A. Robert, V. Greiff, E. Smorodina, P. Rawat, R. Akbar, K. Dreczkowski, R. Tutunov, D. Bou-Ammar, J. Wang, A. Storkey, H. Bou-Ammar, “Toward real-world automated antibody design with combinatorial Bayesian optimization” Cell Rep. Methods 2023, 3, 100374.

[26] S. Stanton, W. J. Maddox, N. Gruver, P. Maffettone, E. Delaney, P. Greenside, A. G. Wilson, “Accelerating Bayesian Optimization for Biological Sequence Design with Denoising Autoencoders”, arXiv 2022, 2203.12742

[27] B. L. Hie, V. R. Shanker, D. Xu, T. U. J. Bruun, P. A. Weidenbacher, S. Tang, W. Wu, J. E. Pak, P. S. Kim, “Efficient evolution of human antibodies from general protein language models” Nat. Biotechnol. 2024, 42, 275–283.

[28] K. Jiang, Z. Yan, M. Di Bernardo, S. R. Sgrizzi, L. Villiger, A. Kayabolen, B. J. Kim, J. K. Carscadden, M. Hiraizumi, H. Nishimasu, J. S. Gootenberg, O. O. Abudayyeh, “Rapid in silico directed evolution by a protein language model with EVOLVEpro” Science 2025, 387, eadr6006.

[29] J. Yang, R. G. Lal, J. C. Bowden, R. Astudillo, M. A. Hameedi, S. Kaur, M. Hill, Y. Yue, F. H. Arnold, “Active learning-assisted directed evolution” Nat. Commun. 2025, 16, 714.

[30] Angermueller, C., Dohan, D., Belanger, D., Deshpande, R., Murphy, K., & Colwell, L. (2020). Model-based reinforcement learning for biological sequence design. International Conference on Learning Representations (ICLR 2020). https://openreview.net/forum?id=HklxbgBKvr

