## Supplementary Information for "BioPipelines: Accessible Computational Protein and Ligand Design for Chemical Biologists"

#### Supplementary figures

```
#resources, PDB, ProteinMPNN and AlphaFold steps (see Figure 1).
conf_change = ConformationalChange(reference_structures = ubiquitin,
                                   target_structures = folded)
filtered_sequences = Panda(tables=[folded.tables.confidence,
                                   conf_change.tables.changes],
                           operations=[Panda.merge(),
                                       Panda.filter("RMSD < 1.5 and plddt > 80")],
                           pool=sequences)
dna = DNAEncoder(sequences=filtered_sequences, organism="EC")
```

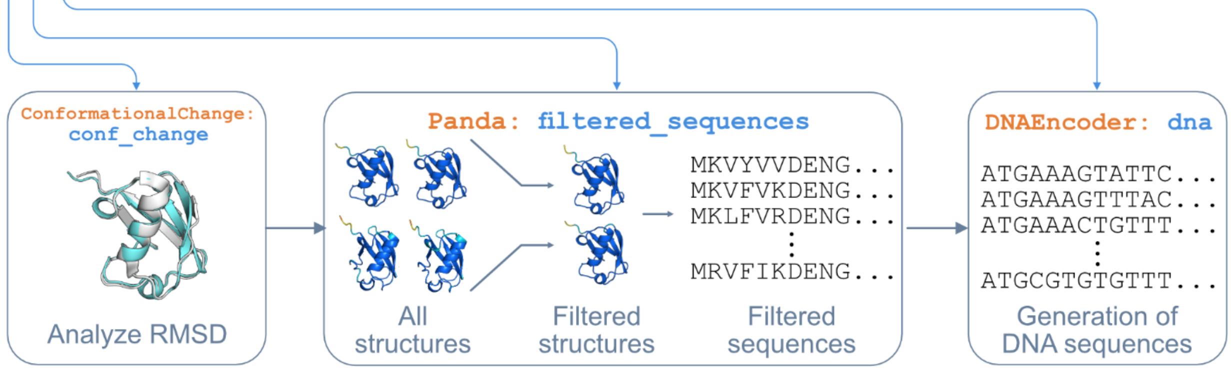

**Figure S1.** The tool ConformationalChange can be used to measure the RMSD between two sets of proteins matched by ID, like apo and holo Boltz folds, or, as done here, between a set of target proteins and a single reference structure. The output is a table containing per-structure RMSD data, which is here matched to the AlphaFold confidence by Panda. The resulting combined data is filtered to only include rows with RMSD < 1.5 Å and plddt > 80, and to extract pool elements, in this case protein sequences, matching the ids that pass the filter.

```
conf_change = ConformationalChange(reference_structures = kinase,
                                   target_structures = refolded,
                                   selection = backbones.tables.structures.fixed )
top3 = Panda(tables=[refolded.tables.confidence,
                     conf_change.tables.changes],
             operations=[Panda.merge(),
                         Panda.filter("RMSD < 1.5")],
                         Panda.sort("plddt",
                                    ascending = False)],
                         Panda.head(3)],
             pool=refolded)
PyMOL(PyMOL.Load(top3),
      PyMOL.ColorAF(),
      PyMOL.Color("white",selection=backbones.tables.structures.fixed)
```

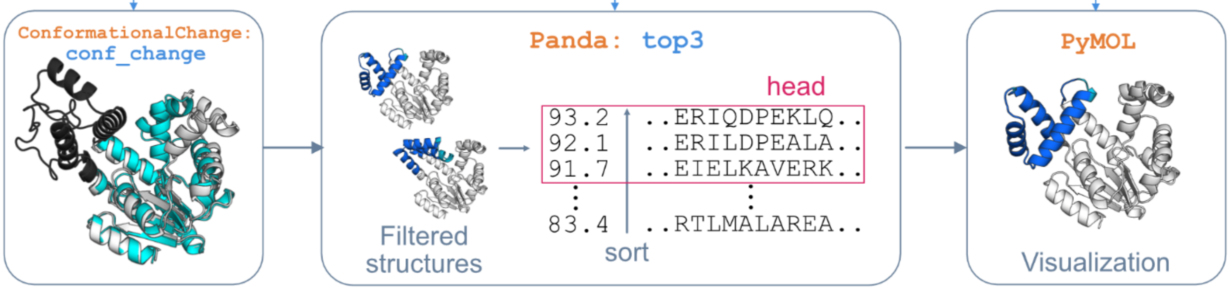

**Figure S2**. Structures whose id pass the series of pandas operations are transferred from the pool to top3. A PyMOL session is created, already displaying the portions of interest.

```
with Pipeline(project="TrpR", job="LibraryScreen"):
       # Resources, Sequence, and Boltz2 identical to Figure3
       SMILES_library = {"candidate": r"<R1>CC1=CNC2=CC=CC=C21",
                         "R1": ["<amine>", "<amino acid>", "<acid>"],
                         "amine": r"[C@@H](N)(C(C)(C)C)",
                         "amino acid": r"[C@@H](N)(C(=O)O)",
                         "acid": r"[C@@H](C(C)(C)C)(C(=O)O)"}
       library = CompoundLibrary(SMILES_library,
                                 generate_images = True)
```

CompoundLibrary: library

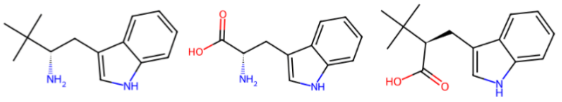

cand0 cand1 cand2

| id | format | smiles | ccd | R1 |
|---|---|---|---|---|
| cand0 | smiles | [C@@H](N)(C(C)(C)C)... | | amine |
| cand1 | smiles | [C@@H](N)(C(=O)O)C... | | amino acid |
| cand2 | smiles | [C@@H](C(C)(C)C)(C(=... | | acid |

**Figure S3**. A compound library can be defined, besides importing a ChemDraw file with R-group tables, also with a SMILES dictionary (which can be loaded from a csv file). We recommend generating compound images for inspection.

```
merged = Panda(tables=[cofolded.tables.affinity,
                       library.tables.compounds],
               operations=[Panda.merge(),
                           Panda.calculate({"aff_uM":"10**affinity_pred_value"})])
Plot(Plot.Scatter(data=merged.tables.result,
                  title="Predicted Affinity by R1",
                  x="R1",
                  y="aff_uM",
                  xlabel="R1 group"
                  ylabel="Predicted Affinity [uM]"))
```

Plot

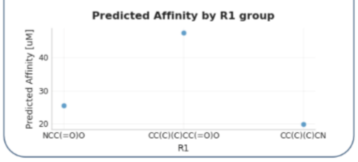

**Figure S4.** The predicted affinity is converted into micromolar units with Panda and plotted against the R1 substituent. When names are not provided for each R1 group in the ChemDraw XML file, the SMILES corresponding to each fragment are displayed.

```
mutants = Mutagenesis(original=fusions,
                      position=Selection(Selection.add(fusions.tables.sequences.L1),
                                         Selection.add(fusions.tables.sequences.L2))
                      mutate_to="LK",
                      combinatorial=True)
# Following steps using sequences from mutants instead of fusions
```

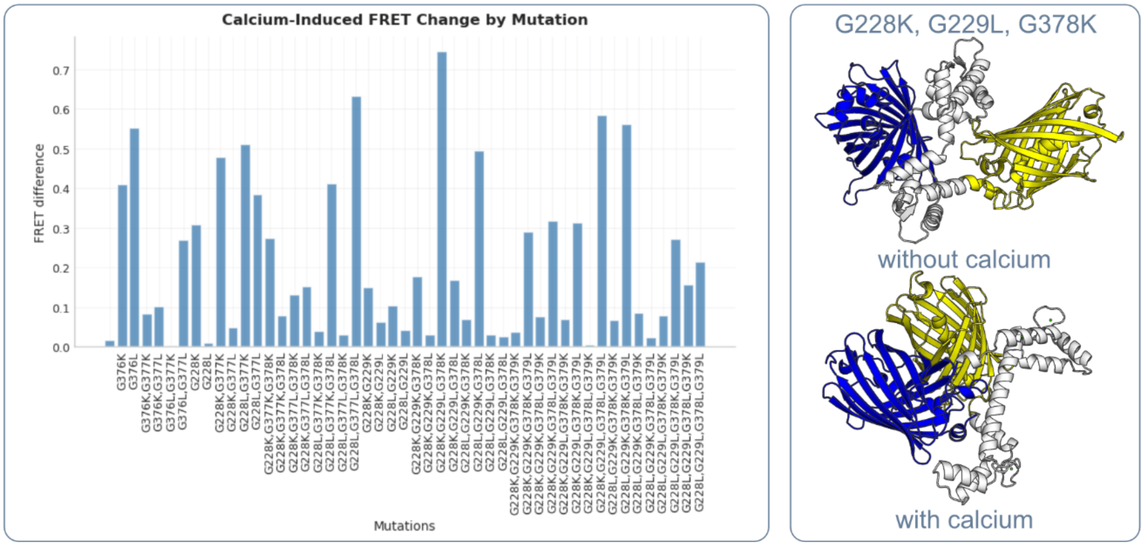

**Figure S5**. Combining linker length and residue screening when designing a FRET sensor. After fusing the sequences with the Fuse tool, here we mutate all the linker positions, which are defined in the sequences table of the Fuse output, populated at runtime, using the Mutagenesis tool. We combine the two residue selections with the Selection tool and feed all the positions to a single instance of Mutagenesis. We chose to mutate to either leucine (L) or lysine (K). By setting the combinatorial mode, we make sure all the resulting sequences have all the residues defined in the selections mutated at once.

```
with Pipeline(project="FRET", job="FRET sensor"):
    dist_apo = Distance(structures = apo,
                        residue = ["66","-173"],
                        metric_name = "FRET_distance_apo")
    dist_holo = Distance(structures = holo,
                         residue = ["66","-173"],
                         metric_name = "FRET_distance_holo")
    R0 = 35.4 # Angstrom
    derived_metrics = {"FRET_E_apo": f"1 / (1 + (FRET_distance_apo / {R0})**6)",
                       "FRET_E_holo": f"1 / (1 + (FRET_distance_holo / {R0})**6)",
                       "delta_FRET": "abs(FRET_E_holo - FRET_E_apo"}
    analysis = Panda(tables=[fusion.tables.sequences,
                             dist_apo.tables.distances,
                             dist_holo.tables.distances],
                     operations=[Panda.merge(),
                                 Panda.calculate(derived_metrics)])
    Plot(Plot.Bar(data=analysis.tables.result,
                  title="FRET efficiency by linker length",
                  x="lengths",
                  y="FRET_E_apo",
                  y_right="FRET_E_holo",
                  xlabel="Linker Lengths"
                  ylabel="FRET apo",
                  ylabel_right_right="FRET holo"))
```

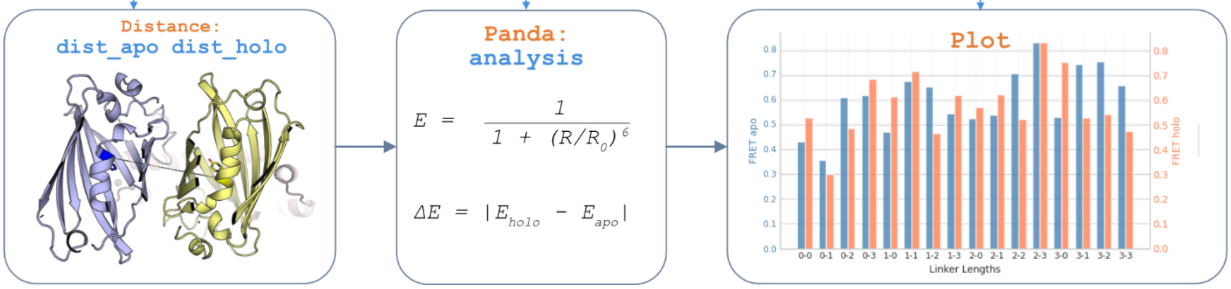

**Figure S6**. Modelling a FRET-based calcium sensor based on calmodulin and two fluorescent proteins. With the Distance tool, we measure the distance between the two fluorophores, positioned at residue 66 in EBFP, and -173 (counting from the C terminus) in EYFP. Further, we use Panda to calculate derived metrics based on the Förster radius of 35.4 Å for the EBFP-EYFP couple, and plot the results against the linker lengths.

```python
with Pipeline(project="Kinase", job="LID_Redesign_Inspection"):
      kinase = PDB("4AKE")
      backbones = RFdiffusion(pdb=kinase),
                              contigs='A1-117/50-70/A161-214',
                              num_designs=10)
      sequences = ProteinMPNN(structures=backbones,
                              num_sequences=2,
                              redesigned=backbones.tables.structures.designed)
sequences
```

**sequences**

STREAMS

| name | format | items | map_table |
|---|---|---|---|
| sequences | csv | 20 | <output_folder>/queries.csv |

sequences

| name | format | items | map_table | files_contain_wildcards |
|---|---|---|---|---|
| sequences | csv | 20 | <output_folder>/queries.csv | False |

| id | gaps | sequence | T | sample | score | global_score | seq_recovery | structures.id |
|---|---|---|---|---|---|---|---|---|
| LID_Redesign_002_6_1 | | MRIILLGAPGAGKGTQAQFIMEKYGIPQISTGDMLRAAVKSGSELGKQAKDIMDAGK... | 0.1 | 1 | 1.0641 | 1.477 | 0.0299 | LID_Redesign_002_6 |
| LID_Redesign_002_6_2 | | MRIILLGAPGAGKGTQAQFIMEKYGIPQISTGDMLRAAVKSGSELGKQAKDIMDAGK... | 0.1 | 2 | 1.0837 | 1.4531 | 0.0299 | LID_Redesign_002_6 |
| ... 16 more ... | | | | | | | | |
| LID_Redesign_002_3_1 | | MRIILLGAPGAGKGTQAQFIMEKYGIPQISTGDMLRAAVKSGSELGKQAKDIMDAGK... | 0.1 | 1 | 1.1609 | 1.5006 | 0.02 | LID_Redesign_002_3 |
| LID_Redesign_002_3_2 | | MRIILLGAPGAGKGTQAQFIMEKYGIPQISTGDMLRAAVKSGSELGKQAKDIMDAGK... | 0.1 | 2 | 1.1604 | 1.5023 | 0.04 | LID_Redesign_002_3 |

TABLES

| name | columns | path |
|---|---|---|
| sequences | id, structures.id, source_pdb, sequence, score, seq_recovery, gaps | <output_folder>/queries.csv |
| missing | id, removed_by, cause | <output_folder>/missing.csv |

```python
with Pipeline(project="AdenylateKinase", job="LID_Redesign_Inspection"):
      backbones = Load("/path/to/002_RFdiffusion")
backbones
```

**backbones**

STREAMS

| name | format | items | map_table |
|---|---|---|---|
| structures | pdb | 10 | <output_folder>/structures_map.csv |

structures

| name | format | items | map_table | files_contain_wildcards |
|---|---|---|---|---|
| structures | pdb | 10 | <output_folder>/structures_map.csv | False |

| id | file | value |
|---|---|---|
| LID_Redesign_002_1 | /shares/locbp.chem.uzh/gquarg/BioPipelines/AdenylateKinas... | |
| LID_Redesign_002_2 | /shares/locbp.chem.uzh/gquarg/BioPipelines/AdenylateKinas... | |
| ... 6 more ... | | |
| LID_Redesign_002_9 | /shares/locbp.chem.uzh/gquarg/BioPipelines/AdenylateKinas... | |
| LID_Redesign_002_10 | /shares/locbp.chem.uzh/gquarg/BioPipelines/AdenylateKinas... | |

TABLES

| name | columns | path |
|---|---|---|
| structures | id, source_id, pdb, fixed, designed, contigs, time, status | <output_folder>/rfdiffusion_results.csv |

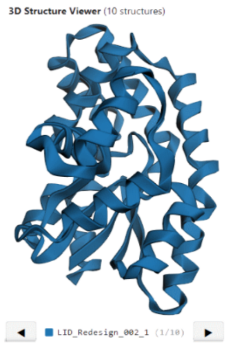

**Figure S7**. Top: Inspection of an intermediate tool's output of a pipeline run in Jupyter. Bottom: Inspection of a tool's output from a previously run pipeline.

**Table S1**. Tools currently implemented in BioPipelines

| Function | # | Tool | Description | Ref. |
|---|---|---|---|---|
| Structure generation | 1 | Rfdiffusion | Protein backbone generation | [1] |
| | 2 | Rfdiffusion All-Atom | Backbone generation, atoms-aware | [2] |
| | 3 | RFdiffusion3 | Protein generation | [3] |
| | 4 | BoltzGen | Protein generation and analysis | [4] |
| Sequence design | 5 | ProteinMPNN | Inverse folding | [5] |
| | 6 | LigandMPNN | Inverse folding, ligand-aware | [6] |
| | 7 | Mutagenesis | Site-directed mutagenesis | |
| | 8 | MutationComposer | Sampling-based sequence generation | |
| | 9 | Fuse | Combinatorial fusing of sequences | |
| | 10 | StitchSequences | Stitching of sequences | |
| | 11 | DNAEncoder | Inverse-translation | [7] |
| | 12 | RBSDesigner | Optimization of the ribosomal binding site | [8] |
| Structure prediction | 13 | AlphaFold2 | Protein folding | [9] |
| | 14 | Boltz2 | Folding of several molecular types | [10] |
| | 15 | Gnina | Molecular docking with deep learning | [11] |
| Basic input types | 16 | Ligand | Ligand loading incl. Pubchem, RCSB | |
| | 17 | PDB | Protein complexes loading incl. RCSB | |
| | 18 | RCSB | Data loading from RCSB using Query API | |
| | 19 | CompoundLibrary | SMILES library expansion | |
| | 20 | Sequence | Protein, DNA, RNA sequences loading. | |
| | 21 | Table | Table loading from files. | |
| Structural analysis | 22 | Distance | Distance calculations (atoms, residues) | |
| | 23 | Angle | Angle and dihedral calculations | |
| | 24 | Contacts | Contacts between ligands and proteins | |
| | 25 | ConformationalChange | Measurement of structural changes | |
| | 26 | PoseChange | Measurement of ligand pose changes | |
| | 27 | PoseBusters | Conformational plausibility | [12] |
| Dynamics | 28 | CABSflex | Coarse-Grained Monte Carlo | [13] |
| Sequence statistics | 29 | MutationProfiler | Profiling of amino acid frequencies | |
| | 30 | SequenceMetricCorrelation | Correlation between a sequence and a metric | |
| | 31 | BayesianAdjuster | Frequency adjustment based on correlations | |
| MSAs | 32 | MMseqs2,MMseqs2Server | Multiple sequence alignment (uniref30, GPU) | [14] |
| Table operations | 33 | Panda | Table operations and stream filtering | |
| | 34 | ExtractMetrics | Metric extraction from several tables | |
| Visualization | 35 | PyMOL | Sessions with aligned and colored structures | |
| | 36 | Plot | Various kinds of plots. | |
| Utility | 37 | LoadOutput | Loading of previously run tools | |

#### Architecture

##### 1. Multimodal execution with a simple syntax

BioPipelines was developed around a desired clear, simple syntax according to which a tool is configured and run by simply writing its name together with its own parameters, and the output can be passed to subsequent tools without acting on it. This result was achieved through a combination of architectural choices and Python's programming language innate features. Namely, a BioPipelines workflow is defined within a Python context manager *with*, through a general Pipeline class, such that tools instantiated inside the context are automatically registered with the pipeline. The autoregistration mechanism is implemented once and for all in the __*new*__ method, as defined in the abstract class BaseConfig, from which all tool configurators inherit. The Pipeline object is responsible for coordinating the tools, that is, chaining their execution, and defining the broad filesystem structure. In general, each pipeline produces a self-contained directory (**FigureS8**). During the configuration phase, the framework predicts all output file paths and generates self-contained bash execution scripts, which are stored in the runtime directory for potential inspection. Each tool's output resides in a numbered, tool-named subdirectory containing the expected output as well as its execution log.

```
<biopipelines_output>/<project>/<job>_<NNN>/
├── RunTime/                  # Execution scripts
│   ├── pipeline.sh
│   ├── 001_<tool>.sh
│   └── 002_<tool>.sh
├── Logs/                     # Execution logs
│   ├── 001_<tool>.log
│   └── 002_<tool>.log
├── ToolOutputs/              # Tool output predictions
│   ├── 001_<tool>.json
│   └── 002_<tool>.json
├── 001_<Tool>/               # Tool outputs
│   ├── .expected_outputs.json
│   ├── .log
│   └── various output (PDBs, CSVs, ...)
└── 002_<Tool>/
... └── ...
```

**Figure S8**. Self-contained output organization of a pipeline's output.

The context manager allows for the pipeline to pick up not only tool configurations as they are added, but also intermediate directives for the overall pipeline script being generated. For example, resources, dependencies, or simple folder name manipulation with a suffix can be appended in the same syntax and are applied either to the slurm script or to the file system structure. A dedicated submission script accessible after package installation via command line interface (CLI) *bp-submit <pipeline.py>* collects the coordinator scripts generated during the configuration and submits them while incorporating potential intra-pipeline dependencies, thus allowing the user to define multi-job pipelines within the same framework (**FigureS9**). Furthermore, with its simple on-the-fly parameter, automatically set to True when a notebook is detected, each tool script *<N>_<tool>.sh* is run as soon as it is generated.

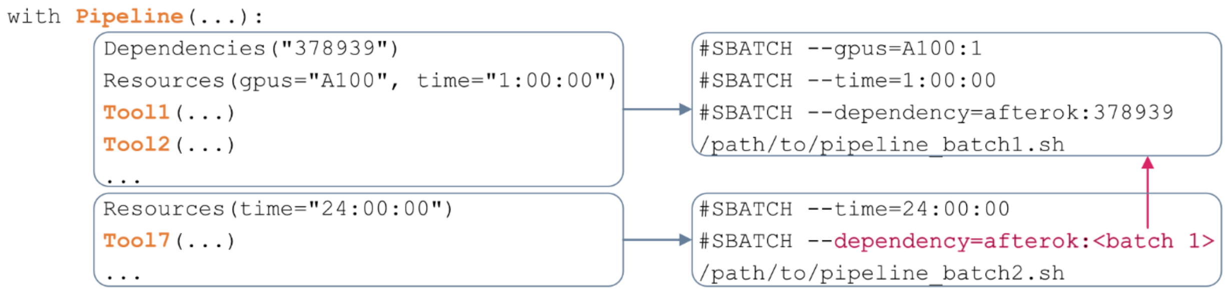

**Figure S9**.Keywords *Dependencies* and *Resources* are interpreted by the biopipelines submit script. When requesting two (or more) resources within the same pipeline, the script will generate and submit two (or more) SLURM batches, each one dependent on the previous one.

#### 2. Standardized interfacing of tools

There are three main challenges in chaining tools within a pipeline: (i) tools organize their outputs arbitrarily; (ii) the parenting relationship between input and output data is possibly lost during execution; and (iii) tools produce and consume different data types for same entity types, for example, common PDB or CIF for structures, CSV or fasta for sequences, and so on. BioPipelines addresses these challenges with the introduction of a StandardizedOutput class exposing a collection of any of two complementary data containers: streams and tables (**Figure S10**).

```python
@dataclass
class Datastream:
        conf_change = str              # "structures", "sequences", ...
        ids = List[str]                # Unique identifiers in pattern form
        map_table = str                # CSV path mapping IDs to files, values, metadata
        files = List[str]              # File paths, in pattern form
        format = str                   # "pdb", "cif", ...
        metadata = List[str, Any]      # Additional stream information

@dataclass
class TableInfo:
        name= str                      # "confidence", "affinity", ...
        path = str                     # Path to CSV file
        columns = List[str]            # Expected column names
        description = str              # Human-readable description
```

**Figure S10**. Public attributes of the DataStream and TableInfo class.

*Streams* (DataStream) represent ID-tracked collections of entities that are produced by and flow between tools. In particular, each stream associates unique identifiers with either files (*.pdb*, *.cif*, …) or values (*"MHHHHHHLTKW…", "ATP", …*), and records the data format. This source tool-agnostic definition of the output implies that, for example, ProteinMPNN can operate on the structures produced by RFdiffusion, PDB, or AlphaFold, without distinction, because all it receives is a DataStream identically formatted.

Stream objects are constructed deterministically during the configuration phase by the tool configurator classes, based on input streams from previous tools and tool-specific parameters. This happens as soon as the tools are added to the pipeline, before the files or values exist. The parent-tool tracking, necessary to compare or combine data from different tools, happens by following a series of simple conventions for ID generation, based on a tool's input vs output relationship:

- 1 entity → 1 entity: output IDs are identical to input IDs. *Example: AlphaFold: 1 sequence → 1 best-ranked model*
- 1 entity → N entities: output IDs are obtained by appending an underscore and a suffix, determined by tool-specific parameters. *Example: ProteinMPNN: 1 structure → N sequences where N is num_sequences. 3KU8 → 3KU8_<1..N>.*
- J entities x K entities x … → N entities: output IDs are combinations of input ones, separated by a + character, and map_table will contain the source IDs as provenance. *Example: Boltz2: [prot1, prot2, …] x [lig1,lig2,…] → [prot1+lig1,prot1+lig2,…].*

The Biopipelines Python package provides standardized functions to handle streams and to match data derived by upstream or downstream tools based on these conventions, and the ReMap tool for reassigning IDs, for example, to combine outputs coming from different pipelines.

*Tables* (TableInfo) represent tabular metadata associated with streams. They describe CSV files and predict column names as well. Tables always present a first column called *id* for matching data to streams, but they are not ID-tracked to allow easier operations, with the tool Panda performing both table operations and filtering of streams based on table data.

When a tool registers itself within an active pipeline context, it returns a *StandardizedOutput* object rather than the tool instance itself. This design is consistent with the principle that once a tool's configuration is concluded, i.e., when the constructor arguments are set, only the predicted output is needed by downstream tools. Further, the *StandardizedOutput* is serializable, such that each tool saves its own predicted output in its output folder as a JSON file, and the file itself can be loaded with the Load tool to obtain an object that behaves identically to the original input. This serialization enables workflows to be resumed or extended across sections.

Finally, we address the problem of different formats with some basic input entity types that ensure representation coherence across the framework. These types are strictly associated with standard streams.  The general practice is that whenever a tool accepts, for example, a protein sequence, it does so by receiving the output of a Sequence object instantiated with the given sequence (**Figure S11**). By enforcing this behavior, tools are guaranteed to operate on tool output streams in the same way as they operate on simple inputs. To extend the framework to new basic types, all that is there to be done is to define a new stream name. The dot access notation holds for any stream name.

```
with Pipeline(...):
        # correct: receives sequences stream from Sequence as from any other tool
        Tool(proteins=Sequence("MHHHHHHGTRK..."))
        # wrong: non-standardized input, no id tracking
        Tool(proteins="MHHHHHHGTRK...))
```

**Figure S11**. Convention for passing stream types as arguments to tool constructors.

#### Tool integration

All tools inherit from BaseConfig and implement four core methods for validation of input parameters, configuration of the output folder with respect to the active pipeline, generation of an execution script, and prediction of the output structure, plus an optional installation method (**Figure S12**).

```
class BaseConfig(ABC):
        TOOL_NAME: str
        def validate_params(self):
        # checks parameters are correct, e.g., num_sequences > 0
        def configure_inputs(self, pipeline_folders: Dict[str,str]):
        # receives folders structure from the active pipeline
        def generate_script(self):
        # generates a bash script for tool execution
        def get_output_files(self):
        # predicts DataStream and TableInfo outputs
        def _install_script(self,folders, env_manager, force_reinstall, **kwargs):
        # produces a script to install the tool

class ProteinMPNN(BaseConfig):
        ...
        def generate_script(self):
         """Generate ProteinMPNN execution script."""
         script_content = "#!/bin/bash\n"
         script_content += "# ProteinMPNN execution script\n"
         script_content += self.generate_completion_check_header()
         script_content += self.activate_environment()
         script_content += self._generate_script_prepare_inputs()
         script_content += self._generate_script_run_proteinmpnn()
         script_content += self._generate_script_create_table()
         script_content += self.generate_completion_check_footer()
         return script_content
```

**Figure S12**. Base configurator class and example of script generation.

Each generated bash script follows a standard five-step structure:

1. Completion check: Skip execution if a completion marker already exists, enabling idempotent re-execution after partial failures.
2. Environment activation: Activate the tool's conda environment as specified in config.yaml.
3. Tool execution: Run the tool's command-line interface with the configured parameters.
4. Output parsing: Execute a helper script (pipe_*.py from HelpScripts/) to parse tool-specific outputs into standardized CSV tables.
5. Output verification: Check that all expected outputs exist and create a completion marker on success.

Steps (1) and (5) are handled by biopipelines via a shared script since every tool's own output is defined as a JSON-serialized object, and also environment activation based on machine configuration is handled by the package.

**Configuration**

The config.yaml file at the repository root specifies cluster-specific settings: output paths, conda environment names for each tool, SLURM partition and account information, and GPU type mappings (**Figure S13**). This single configuration file is the only machine-specific component; all pipeline scripts are portable across installations. A Colab-specific configuration (colab.yaml) is present since Google Colab doesn't provide conda. BioPipelines auto-detects Colab environment and loads colab.yaml, which sets env_manager: "pip" and disables environment activation.

```yaml
folders:
  base:
    home: "/home/<username>"
    data: "<home>/data"                    # where tool repositories are installed
    group: "/shares/your_group"            # shared group storage
    user: "<group>/<username>"
    biopipelines: "<cwd>"                  # auto-set to repository root
    biopipelines_output: "<user>/BioPipelines"

  infrastructure:
    HelpScripts: "<biopipelines>/HelpScripts"
    PDBs: "<biopipelines>/PDBs"
    Ligands: "<biopipelines>/Ligands"
    Sequences: "<biopipelines>/Sequences"
    Tables: "<biopipelines>/Tables"
    cache: "<group>/cache"
    containers: "<group>/containers"

  # Tool installation paths
  repositories:
    RFdiffusion: "<data>/RFdiffusion"
    ProteinMPNN: "<data>/ProteinMPNN"
    LigandMPNN: "<data>/LigandMPNN"
    # ...

environments:
  RFdiffusion: "SE3nv"
  ProteinMPNN: "SE3nv"
  LigandMPNN: "ligandmpnn_env"
  # ...

cluster:
  emails:
    your_username: ""
  env_manager: "mamba"            # "mamba" or "conda"
  container_executor: "apptainer" # "apptainer" or "singularity"
  slurm_modules:                  # modules loaded in SLURM scripts
    - "miniforge3"
    - "apptainer"
```

**Figure S13**. Machine configuration via yaml file.

#### References

[1] J. L. Watson, D. Juergens, N. R. Bennett, B. L. Trippe, J. Yim, H. E. Eisenach, W. Ahern, A. J. Borst, R. J. Ragotte, L. F. Milles, B. I. M. Wicky, N. Hanikel, S. J. Pellock, A. Courbet, W. Sheffler, J. Wang, P. Venkatesh, I. Sappington, S. V. Torres, A. Lauko, V. De Bortoli, E. Mathieu, S. Ovchinnikov, R. Barzilay, T. S. Jaakkola, F. DiMaio, M. Baek, D. Baker, "De novo design of protein structure and function with RFdiffusion" *Nature* **2023**, *620*, 1089–1100.

[2] R. Krishna, J. Wang, W. Ahern, P. Sturmfels, I. Kalvet, G. R. Lee, F. S. Morey-Burrows, I. R. Humphreys, R. McHugh, D. Vafeados, X. Li, A. Sutherland, A. Hitchcock, C. N. Hunter, M. Baek, F. DiMaio, "Generalized Biomolecular Modeling and Design with RoseTTAFold All-Atom", *Science*, **2024**, Vol 384, Issue 6693.

[3] J. Butcher, R. Krishna, R. Mitra, R. I. Brent, Y. Li, N. Corley, P. Kim, J. Funk, S. Mathis, S. Salike, A. Muraishi, H. Eisenach, T. R. Thompson, J. Chen, Y. Politanska, E. Sehgal, B. Coventry, O. Zhang, B. Qiang, K. Didi, M. Kazman, F. DiMaio, D. Baker, "De novo design of all-atom biomolecular interactions with RFdiffusion3" *bioRxiv* **2025**, 2025.09.18.676967.

[4] H. Stark, F. Faltings, M. Choi, Y. Xie, E. Hur, T. O'Donnell, A. Bushuiev, T. Uçar, S. Passaro, W. Mao, M. Reveiz, R. Bushuiev, T. Pluskal, J. Sivic, K. Kreis, A. Vahdat, S. Ray, J. T. Goldstein, A. Savinov, J. A. Hambalek, A. Gupta, D. A. Taquiri-Diaz, Y. Zhang, A. K. Hatstat, A. Arada, N. H. Kim, E. Tackie-Yarboi, D. Boselli, L. Schnaider, C. C. Liu, G.-W. Li, D. Hnisz, D. M. Sabatini, W. F. DeGrado, J. Wohlwend, G. Corso, R. Barzilay, T. Jaakkola, **2025**, DOI: 10.1101/2025.11.20.689494.

[5] J. Dauparas, I. Anishchenko, N. Bennett, H. Bai, R. J. Ragotte, L. F. Milles, B. I. M. Wicky, A. Courbet, R. J. De Haas, N. Bethel, P. J. Y. Leung, T. F. Huddy, S. Pellock, D. Tischer, F. Chan, B. Koepnick, H. Nguyen, A. Kang, B. Sankaran, A. K. Bera, N. P. King, D. Baker, "Robust deep learning–based protein sequence design using ProteinMPNN" *Science* **2022**, *378*, 49–56.

[6] J. Dauparas, G. R. Lee, R. Pecoraro, L. An, I. Anishchenko, C. Glasscock, D. Baker, "Atomic context-conditioned protein sequence design using LigandMPNN" *Nat. Methods* **2025**, *22*, 717–723.

[7] A. Alexaki, J. Kames, D. D. Holcomb, J. Athey, L. V. Santana-Quintero, P. V. N. Lam, N. Hamasaki-Katagiri, E. Osipova, V. Simonyan, H. Bar, A. A. Komar, C. Kimchi-Sarfaty, "Codon and Codon-Pair Usage Tables (CoCoPUTs): Facilitating Genetic Variation Analyses and Recombinant Gene Design" *J. Mol. Biol.* **2019**, *431*, 2434–2441.

[8] H. M. Salis, E. A. Mirsky, C. A. Voigt, “Automated design of synthetic ribosome binding sites to control protein expression” *Nat. Biotechnol.* **2009**, *27*, 946–950.

[9] J. Jumper, R. Evans, A. Pritzel, T. Green, M. Figurnov, O. Ronneberger, K. Tunyasuvunakool, R. Bates, A. Žídek, A. Potapenko, A. Bridgland, C. Meyer, S. A. A. Kohl, A. J. Ballard, A. Cowie, B. Romera-Paredes, S. Nikolov, R. Jain, J. Adler, T. Back, S. Petersen, D. Reiman, E. Clancy, M. Zielinski, M. Steinegger, M. Pacholska, T. Berghammer, S. Bodenstein, D. Silver, O. Vinyals, A. W. Senior, K. Kavukcuoglu, P. Kohli, D. Hassabis, “Highly accurate protein structure prediction with AlphaFold” *Nature* **2021**, *596*, 583–589.

[10] S. Passaro, G. Corso, J. Wohlwend, M. Reveiz, S. Thaler, V. R. Somnath, N. Getz, T. Portnoi, J. Roy, H. Stark, D. Kwabi-Addo, D. Beaini, T. Jaakkola, R. Barzilay, “Boltz-2: Towards accurate and efficient binding affinity prediction” *bioRxiv* **2025**, 2025.06.14.659707.

[11] A. T. McNutt, P. Francoeur, R. Aggarwal, T. Masuda, R. Meli, M. Ragoza, J. Sunseri, D. R. Koes, “GNINA 1.0: molecular docking with deep learning” *J. Cheminformatics* **2021**, *13*, 43.

[12] M. Buttenschoen, G. M. Morris, C. M. Deane, “PoseBusters: AI-based docking methods fail to generate physically valid poses or generalise to novel sequences” *Chem. Sci.* **2024**, *15*, 3130–3139.

[13] A. Kuriata, A. M. Gierut, T. Oleniecki, M. P. Ciemny, A. Kolinski, M. Kurcinski, S. Kmiecik, “CABS-flex 2.0: a web server for fast simulations of flexibility of protein structures” *Nucleic Acids Res.* **2018**, *46*, W338–W343.

[14] M. Steinegger, J. Söding, “MMseqs2 enables sensitive protein sequence searching for the analysis of massive data sets” *Nat. Biotechnol.* **2017**, *35*, 1026–1028.

**Appendix A**

**Creation of PoseBusters tool with Claude Code Opus 4.6**

The following is the transcription of a conversation with state-of-the-art reasoning model Claude Code Opus 4.6 to implement the PoseBusters tool. We have highlighted in red the input from the user, and in blue the output from Claude Code.

USER ❯ Enact Docs/LLM_prompt.txt. Implement the tool PoseBusters (https://github.com/maabuu/posebusters), conforming to the existing tool standards.

CLAUDE > ● I'll start by reading the LLM prompt document and exploring the codebase to understand the existing tool standards.

● Read 1 file (ctrl+o to expand)

● Explore(Explore codebase structure and patterns)

Done (29 tool uses · 82.2k tokens · 1m 45s)

(ctrl+o to expand)

● Now let me read the referenced docs and explore the PoseBusters tool to understand what it does.

● Read 3 files (ctrl+o to expand)

● Good. Now let me look at a concrete tool implementation closely, and explore PoseBusters to understand what it does.

● Read 1 file (ctrl+o to expand)

● Fetch(https://github.com/maabuu/posebusters)

Received 263.3KB (200 OK)

● Let me get more details on PoseBusters and examine the existing tool patterns more closely.

● Fetch(https://github.com/maabuu/posebusters/blob/main/README.md)

Received 197.3KB (200 OK)

● Read 1 file (ctrl+o to expand)

● Fetch(https://posebusters.readthedocs.io)

Received 23KB (200 OK)

● Read 1 file (ctrl+o to expand)

● Now let me look at an existing HelpScript and the config.yaml to understand the environment setup pattern, and also check if there's a tool similar to PoseBusters (analysis tool working with protein-ligand structures).

● Read 2 files (ctrl+o to expand)

● Now let me check the __init__.py to see how tools are exported, and look at the PoseChange tool since it's closely related.

● Searched for 4 patterns, read 2 files (ctrl+o to expand)

● Let me read the PoseChange tool since it's the most similar tool (ligand pose analysis), and also peek at the colab.yaml to understand pip mode.

● Read 2 files (ctrl+o to expand)

● Now I have a comprehensive understanding of the codebase. Let me now design the implementation plan.

● Plan(Design PoseBusters implementation plan)

Done (15 tool uses · 117.0k tokens · 2m 22s)

(ctrl+o to expand)

● I have a thorough understanding now. Let me check the __init__.py to see the exact registration pattern before writing the plan.

● Read 1 file (ctrl+o to expand)

● Now I have everything I need. Before writing the final plan, I have a couple of clarifying questions.

● User answered Claude's questions:

(i) PoseBusters needs ligands as SDF files, but your structures from Boltz2/Gnina are PDB/CIF with ligands embedded. The HelpScript needs to extract the ligand to SDF. Should the tool support only `dock` mode (ligand + protein, no reference), or also `redock` mode (ligand + protein + reference ligand for RMSD comparison)? → USER > Both dock and redock.

(ii) PoseBusters requires RDKit (it's a dependency). Should the tool use a dedicated `posebusters` conda environment, or install into the existing `ProteinEnv` (which already has RDKit)? → USER > Dedicated posebusters env (Recommended).

● Good. Now let me write the final plan.

● Updated plan

/plan to preview

***PoseBusters Tool Implementation Plan***

***Context***

*PoseBusters validates computationally generated molecule poses (bond lengths, angles, steric clashes, protein interactions, stereochemistry, etc.). Adding it as a BioPipelines tool enables automated quality validation of docking/prediction outputs from tools like Boltz2 and Gnina.*

Supports both dock mode (ligand + protein) and redock mode (+ reference ligand for RMSD comparison).

**Files to Create**

**1**. biopipelines/posebusters_tool.py — Tool config class

File named posebusters_tool.py to avoid collision with the posebusters pip package.

Pattern: Follow biopipelines/contacts.py exactly.

class PoseBusters(BaseConfig):

TOOL_NAME = "PoseBusters"

- Constructor parameters:

- structures: Union[DataStream, StandardizedOutput] — protein-ligand complexes (PDB/CIF)

- ligand: str — 3-letter residue code for the ligand (e.g., "LIG", "ATP")

- reference_ligand: Union[DataStream, StandardizedOutput, None] = None — for redock mode, reference ligand structure (SDF or PDB with ligand)

- reference_ligand_code: Optional[str] = None — residue code in reference structure (defaults to ligand)

- mode: str = "dock" — "dock" or "redock" (auto-set to "redock" if reference_ligand provided)

- id_map: Dict[str, str] = {"*": "*_<S>"} — standard ID mapping

- Lazy paths:

- analysis_csv → <output_folder>/posebusters.csv

- config_file → <output_folder>/posebusters_config.json

- structures_ds_json → <output_folder>/structures.json

- posebusters_py → <HelpScripts>/pipe_posebusters.py

- _install_script: Creates dedicated posebusters conda env:

- conda/mamba: create -n posebusters python=3.10 -y && pip install posebusters

- pip mode: pip install posebusters

- Skip if already installed (unless force_reinstall)

- validate_params: structures non-empty, ligand is 1-3 char string, mode is "dock"/"redock", redock requires reference_ligand.
- generate_script: Standard pattern — serialize DataStream JSON + config JSON, call python pipe_posebusters.py --config <config.json>.

Config JSON written at config time:

```
{
  "structures_json": "<path>",
  "ligand_name": "LIG",
```

*"mode": "dock",*

*"reference_pdb": null,*

*"reference_ligand_code": null,*

*"id_map": {"*": "*_<S>"},*

*"output_csv": "<path>"*

*}*

*For redock mode, reference_pdb is set to the reference structure file path and reference_ligand_code to the ligand code in the reference.*

- *get_output_files: Returns table "posebusters" with columns dynamically listed (id, source_structure, plus PoseBusters boolean check columns, plus all_pass).*

***2**. HelpScripts/pipe_posebusters.py — Runtime script*

*Pattern: Follow HelpScripts/pipe_contacts.py.*

*Logic flow:*

*1. Parse --config, load JSON*

*2. Load structures DataStream via biopipelines_io.load_datastream() + iterate_files()*

*3. For each structure:*

*a. Extract ligand to SDF — parse PDB/CIF, isolate HETATM records matching ligand code, use RDKit MolFromPDBBlock + rdDetermineBonds.DetermineBonds() for bond perception, write SDF via Chem.SDWriter*

*b. Extract protein to PDB — write all ATOM lines + non-ligand HETATM to a clean PDB*

*c. For redock mode: extract reference ligand from reference structure similarly*

*d. Run PoseBusters via Python API:*

*PoseBusters(config=mode).bust(mol_pred=ligand.sdf, mol_cond=protein.pdb*
*[, mol_true=ref.sdf])*

*e. Collect boolean results dict from the returned DataFrame*

*4. Assemble results into a pandas DataFrame*

*5. Add all_pass column (AND of all boolean check columns)*

*6. Save to output_csv*

*Key implementation details for ligand extraction:*

*def extract_ligand_and_protein(structure_path, ligand_name, output_dir, structure_id):*

"""Extract ligand to SDF and protein to PDB from a PDB/CIF structure."""
ext = os.path.splitext(structure_path)[1].lower()
### Read all lines, separate ligand HETATM from protein ATOM/HETATM
### For CIF: use gemmi to convert to PDB first, then process
### Ligand: RDKit MolFromPDBBlock on isolated HETATM block
### If bonds not assigned (predicted structures lack CONECT):
#   use rdDetermineBonds.DetermineBonds(mol)
### Write to SDF
### Protein: everything except ligand residue, write to PDB
return ligand_sdf_path, protein_pdb_path

CIF handling: Use gemmi (PoseBusters dependency) to read CIF and convert/extract atoms.
gemmi.read_structure(path) works for both PDB and CIF.

Multi-ligand: If multiple copies of the same ligand exist, run
PoseBusters on each copy, suffix the ID with _lig1, _lig2, etc.

**Files to Modify**

**3**. biopipelines/__init__.py

Add import and __all__ entry:

from .posebusters_tool import PoseBusters as PoseBustersTool

Add 'PoseBustersTool' to __all__.

4. config.yaml

Add under environments:

PoseBusters: "posebusters"

5. colab.yaml

No changes needed (pip mode handled by _install_script).

**Reusable Utilities**

- HelpScripts/biopipelines_io.py: load_datastream(), iterate_files() — for loading structures at runtime
- HelpScripts/id_map_utils.py: map_table_ids_to_ids() — for ID mapping
- biopipelines/base_config.py: BaseConfig, StandardizedOutput, TableInfo, Path
- biopipelines/datastream.py: DataStream

**Example Usage**

from biopipelines.pipeline import *
from biopipelines.posebusters_tool import PoseBusters
with Pipeline("Validation", "PoseBusters-Check"):

```
Resources(gpu="A100", time="2:00:00")
PoseBusters.install()
protein = PDB("1ABC")
ligand = Ligand("ATP", codes="ATP")
holo = Boltz2(sequences=protein, compounds=ligand)
# Dock mode
validation = PoseBusters(structures=holo, ligand="ATP")
# validation.tables.posebusters.all_pass -> column reference
# Redock mode (with reference)
reference = PDB("1ABC")  # crystal structure with ligand
validation = PoseBusters(
    structures=holo,
    ligand="ATP",
    reference_ligand=reference,
    reference_ligand_code="ATP",
    mode="redock"
)
```

***Verification***

*1. Import test: from biopipelines.posebusters_tool import PoseBusters should work without error*

*2. Config-time test: Create a PoseBusters instance within a Pipeline context, verify it produces the config JSON and structures JSON in the output folder*

*3. End-to-end test (on-the-fly in notebook): Run with a known Boltz2 output containing a ligand, verify posebusters.csv is produced with boolean check columns and all_pass*

*4. Redock test: Provide a reference structure, verify RMSD column appears in output*

USER > [Accepted plan with cleared context]

● I'll start by reading the key reference files to understand the patterns, then implement the plan.

● Read 4 files (ctrl+o to expand)

● Now let me also look at the base_config for install_script pattern and the gnina tool for a conda env install example.

● Read 3 files (ctrl+o to expand)

● Now let me look at an install script that creates a conda env (like Boltz2) for the pattern.

● Searched for 1 pattern (ctrl+o to expand)

● Now I have all the patterns. Let me create the files. I'll start with the tool config class and the runtime script in parallel, then modify the init and config files.

● Write(biopipelines\posebusters_tool.py)

Wrote 288 lines to biopipelines\posebusters_tool.py

```
# Authors (2026): G. Quargnali & P. Rivera-Fuentes @ LOCBP (https://www.locbp.com/) University of Zurich Switzerland
#
# Licensed under the MIT License. See LICENSE file in the project root for details.

"""
PoseBusters validation for computationally generated molecule poses.

Validates docking/prediction outputs by checking bond lengths, angles,
steric clashes, protein interactions, stereochemistry, and more.
Supports ``dock`` mode (ligand + protein) and ``redock`` mode
```

… +278 lines (ctrl+o to expand)

● Write(HelpScripts\pipe_posebusters.py)

Wrote 366 lines to HelpScripts\pipe_posebusters.py

```
#!/usr/bin/env python3
# Authors (2026): G. Quargnali & P. Rivera-Fuentes @ LOCBP (https://www.locbp.com/) University of Zurich Switzerland
#
# Licensed under the MIT License. See LICENSE file in the project root for details.

"""
Runtime helper script for PoseBusters validation.

Extracts ligands and proteins from predicted structures, runs PoseBusters
quality checks, and outputs a CSV with boolean pass/fail columns
```

.

… +356 lines (ctrl+o to expand)

● Now update __init__.py and config.yaml:

● Update(biopipelines__init__.py)

Added 1 line

```
44  from .mmseqs2 import MMseqs2, MMseqs2Server
45  from .remap import ReMap
46  from .gnina import Gnina
47 +from .posebusters_tool import PoseBusters as PoseBustersTool
48  from .converters import *
49  from .entities import *
50
```

● Update(biopipelines__init__.py)

Added 1 line

```
111      'MMseqs2Server',
112      'ReMap',
113      'Gnina',
114 +    'PoseBustersTool',
115
116      # Utility functions
117      'pdb_to_jsonl',
```

● Update(config.yaml)

Added 1 line

```
 96    MutationProfiler: "MutationEnv"
 97    SequenceMetricCorrelation: "MutationEnv"
 98    BayesianAdjuster: "MutationEnv"
 99 +  PoseBusters: "posebusters"
100    Contacts: "ProteinEnv"
101    PoseChange: "ProteinEnv"
102    SASA: "ProteinEnv"
```

● Now let me verify the import works:

● Bash(cd "C:\Users\gianl\OneDrive\Desktop\PhD\Coding\GitHub-LOCBP\biopipelines"
&& python -c "from biopipelines.posebusters_tool import PoseBusters;

print('Import OK:…)

Import OK: PoseBusters

● Bash(cd "C:\Users\gianl\OneDrive\Desktop\PhD\Coding\GitHub-LOCBP\biopipelines"
&& python -c "from biopipelines import PoseBustersTool; print('Module
import OK:', Pos…)

Module import OK: PoseBusters

● All done. Here's a summary of what was implemented:

Files created:

1. biopipelines/posebusters_tool.py — Tool config class following contacts.py pattern:
 - PoseBusters(BaseConfig) with TOOL_NAME = "PoseBusters"
 - Constructor takes structures, ligand, reference_ligand, reference_ligand_code, mode, id_map
 - Auto-sets mode to "redock" when reference_ligand is provided
 - _install_script creates a dedicated posebusters conda env (Python 3.10)
 - validate_params checks structures non-empty, ligand 1-3 chars, mode validity, redock requires reference
 - generate_script serializes DataStream JSON + config JSON, calls pipe_posebusters.py
 - get_output_files returns posebusters table with boolean check columns + all_pass
2. HelpScripts/pipe_posebusters.py — Runtime script following pipe_contacts.py pattern:
 - Extracts ligands to SDF using gemmi (PDB/CIF) + RDKit (MolFromPDBBlock + DetermineBonds)
 - Extracts protein to clean PDB (everything except ligand)
 - Handles multi-ligand copies (suffixed _lig1, _lig2, etc.)
 - Runs PoseBusters Python API: PoseBusters(config=mode).bust(mol_pred, mol_cond, [mol_true])
 - Computes all_pass column (AND of all boolean checks)
 - Supports redock mode with reference ligand extraction

Files modified:

3. biopipelines/__init__.py — Added from .posebusters_tool import PoseBusters as PoseBustersTool and 'PoseBustersTool' to __all__
4. config.yaml — Added PoseBusters: "posebusters" under environments:

Following this first implementation, the tool was first installed by submitting the following script:

```
from biopipelines.pipeline import *
from biopipelines.posebusters import PoseBusters
with Pipeline("Setup", "InstallPoseBusters"):
      Resources(time="1:00:00", memory="16GB")
      PoseBusters.install()
```

The functionality of the tool was tested by loading some Boltz2 results in Jupyter:

```
from biopipelines.pipeline import *
from biopipelines.load import Load
from biopipelines.posebusters import PoseBusters
with Pipeline("Test", "PoseBusters"):
      boltz_predictions = Load("/path/to/Boltz2 output")
      posebusters = PoseBusters(structures=boltz_predictions,
                                ligand="LIG")
posebusters
```

From which it was found that the package gemmi is required. The installer script was readjusted to include it, the environment was updated, and the same test was run again in Jupyter to obtain the desired analysis (some columns not visible):

**posebusters**

TABLES

| name | columns | path |
|---|---|---|
| posebusters | id, source_structure, mol_pred_loaded, mol_cond_loaded, sanitization, all_atoms_connected, bond_lengths, bond_angles, internal_steric_clash, aromatic_ring_flatness, double_bond_flatness, internal_energy, volume_overlap_with_protein, minimum_distance_to_protein, all_pass | <output_folder>/posebusters.csv |

posebusters

| name | path | columns | description | count |
|---|---|---|---|---|
| posebusters | <output_folder>/posebusters.csv | id, source_structure, mol_pred_loaded, mol_cond_loaded, sanitization, all_atoms_connected, bond_lengths, bond_angles, internal_steric_clash, aromatic_ring_flatness, double_bond_flatness, internal_energy, volume_overlap_with_protein, minimum_distance_to_protein, all_pass | PoseBusters validation (dock mode): LIG | 20 |

| id | source_structure | mol_pred_loaded | mol_cond_loaded | sanitization | inchi_convertible | all_atoms_connected | no_radicals | bond_lengths |
|---|---|---|---|---|---|---|---|---|
| 3QRK_93A_STI | /shares/locbp.chem.uzh/gquarg/BioPipelines/Imatinib/PoseS... | True | True | True | False | True | False | True |
| 3QRK_93C_STI | /shares/locbp.chem.uzh/gquarg/BioPipelines/Imatinib/PoseS... | True | True | True | False | True | False | True |
| 3QRK_93W_STI | /shares/locbp.chem.uzh/gquarg/BioPipelines/Imatinib/PoseS... | True | True | True | False | True | False | True |
| 3QRK_93Y_STI | /shares/locbp.chem.uzh/gquarg/BioPipelines/Imatinib/PoseS... | True | True | True | False | True | False | True |

The whole process took place within 40 min.
